# RNA polymerase II plays an active role in the formation of gene loops through the Rpb4 subunit

**DOI:** 10.1101/602227

**Authors:** Paula Allepuz-Fuster, Michael J. O’Brien, Noelia González-Polo, Bianca Pereira, Zuzer Dhoondia, Athar Ansari, Olga Calvo

## Abstract

Gene loops are formed by the interaction of initiation and termination factors occupying the distal ends of a gene during transcription. RNAPII is believed to affect gene looping indirectly owing to its essential role in transcription. The results presented here, however, demonstrate a direct role of RNAPII in gene looping through the Rpb4 subunit. 3C analysis revealed that gene looping is abolished in the *rpb4*Δ mutant. In contrast to the other looping-defective mutants, *rpb4*Δ cells do not exhibit a transcription termination defect. *RPB4* overexpression, however, rescued the transcription termination and gene looping defect of *sua7-1*, a mutant of TFIIB. Furthermore, *RPB4* overexpression rescued the *ssu72-2* gene looping defect, while *SSU72* overexpression restored the formation of gene loops in *rpb4*Δ cells. Interestingly, the interaction of TFIIB with Ssu72 is compromised in *rpb4*Δ cells. These results suggest that the TFIIB-Ssu72 interaction, which is critical for gene loop formation, is facilitated by Rpb4. We propose that Rpb4 is promoting the transfer of RNAPII from the terminator to the promoter for reinitiation of transcription through TFIIB-Ssu72 mediated gene looping.

## INTRODUCTION

Eukaryotic RNA polymerases (RNAPs) I, II and III are responsible for the synthesis of rRNA, mRNA and tRNA, respectively. RNAPII is the best characterized of all the eukaryotic RNAPs and its structure is highly conserved among eukaryotes [1,2]. During the initiation phase of transcription, RNAPII proceeds through several steps, such as promoter recognition, preinitiation complex (PIC) assembly, open complex formation, initiation and finally promoter escape. The polymerase then makes a transition into the elongation phase. When the polymerase reaches the 3′ end of the gene, termination phase sets in. Following cleavage/polyadenylation of the transcript, polymerase is released from the template and is recycled to the promoter to reinitiate the subsequent round of transcription [3]. In a number of actively transcribing genes, promoter and terminator elements are juxtaposed to form a dynamic structure called a ‘gene loop’. The looped gene architecture is formed by the physical interaction of the initiation and termination factors occupying the distal ends of a gene [4,5]. First described in yeast, gene looping has been demonstrated during transcription of RNAPII-transcribed genes in higher eukaryotes as well [4,6,7]. The general transcription factor TFIIB and Rpb1 carboxy-terminal (Rpb1-CTD) phosphatase Ssu72 [8,9], which is also a component of the CPF termination complex, have been shown to play a crucial role in the formation of gene loops in budding yeast [4,10]. Interestingly, mutations in either TFIIB or Ssu72 that adversely affect gene looping, also had an antagonistic effect on termination of transcription [11,12] as well as transcriptional directionality [13]. Gene looping has also been implicated in reinitiation of transcription, intron-mediated enhancement of transcription and transcription memory [13–15]. Gene looping has been shown to accompany activated transcription only, and consequently is abolished in the absence of transcription activators [14]. Besides activators and TFIIB, TFIIH and Mediator complex also play a critical role in gene loop formation by directly interacting with the CF1 and CPF 3’-end processing/termination factors [12,16]. It is generally believed that RNAPII is not directly involved in formation of gene loops, but is a rather passive player in the process. Since gene looping is a transcription-dependent process, it has been presumed that all mutations that affect transcription will also alter gene looping. Accordingly, gene looping was completely abolished in the *rpb1-1* mutant [17].

The structure of RNAPII is highly conserved among eukaryotes. In all eukaryotes from yeast to humans, RNAPII is composed of 12 subunits. Only *Saccharomyces cerevisiae* RNAPII exists in a 10-subunit core and a dissociable subcomplex formed by Rpb4 and Rpb7 [18,19]. The 12 subunits of RNAPII have been implicated in different functions during the process of transcription. The two largest subunits, Rpb1 and Rpb2, are required for binding to the DNA template. Rpb1 contains a groove for the entry of deoxyribonucleotides. Rpb7 has two RNA binding domains and plays a pivotal role in mRNA decay [20]. Rpb9 has been implicated in start-site selection, but is dispensable for assembly of the 10-subunit enzyme (Hull et al., 1995). While the two largest subunits, Rpb1 and Rpb2, have well-established roles in the initiation of transcription, a few published reports suggest that some of the RNAPII subunits function in the termination of transcription. The CTD of the largest subunit Rpb1 consists of multiple repeats of the heptapeptide sequence (YSPTSPS). The phosphorylation of Ser2 of the CTD is critical for the recruitment of 3’-end processing factors and for termination of transcription [21,22]. In the crystal structure, the Rpb3/Rpb11 heterodimer lies in close proximity of the RNA exit channel [23]. It was therefore proposed that the 3’-end processing factors that contact the nascent RNA also associate with the Rpb3/Rpb11 heterodimer. Accordingly, mutations in subunits Rpb3 *(rpb3E6K)* and Rpb11 (*rpb11E108G*) resulted in the expression of the *URA3* gene located downstream from the terminator region of a reporter gene. This resembles the transcription readthrough phenotype and is indicative of defective termination [24,25]. Intriguingly, mutations in the bacterial counterpart of the Rpb3/Rpb11 heterodimer, the α-subunit homodimer, also led to the readthrough of a termination signal, thereby suggesting a highly conserved role of these subunits in transcription termination [26]. The Rpb4/Rpb7 heterodimer is located near the CTD of the Rpb1 subunit, a position with potential for the interaction with termination factors. In *S. pombe*, the Rpb7 subunit interacts with Seb1, which is the homolog of the Nrd1 termination factor of *S. cerevisiae* [27,28]. Furthermore, the Rpb4/7 dimer is also required for the recruitment of Ssu72 and Fcp1 to the chromatin and the RNAPII [29].

Our data provide evidences for a direct role of RNAPII in the formation of gene loops through the Rpb4 subunit. Although, the absence of Rpb4 does not trigger termination defects, *RPB4* overexpression restores the transcription termination and gene looping deficiencies of the TFIIB mutant *sua7-1*. Similarly, *RPB4* overexpression rescues *ssu72-2* gene looping defects and, *vice versa, SSU72* overexpression restores the formation of gene loops in *rpb4Δ* cells. Interestingly, *in vitro* data demonstrate that Ssu72 interacts with the Rpb4/7 heterodimer through Rpb4. Finally, we found that the TFIIB-Ssu72 interaction is hardly detectable in the absence of Rpb4 and that a complete 12-subunit RNAPII is required for interaction with Ssu72 and TFIIB. We propose that Rpb4 plays a critical role in facilitating the interaction of TFIIB and Ssu72 with each other and with RNAPII during the formation of gene loops, and helps in the transfer of the polymerase from the terminator regions to the promoters to initiate a new round of transcription.

## RESULTS

### Rpb4 is required for the formation of gene loops

Several years ago we demonstrated that the Rpb4/7 heterodimer plays an important role in regulating RNAPII dephosphorylation through Ssu72 and Fcp1 CTD-phosphatases. Ssu72 is not merely a Serine-5/7-CTD phosphatase, but owing to its interaction with the general transcription factor TFIIB, plays a crucial role in gene looping [9,10,17,30]. Furthermore, Ssu72 being a component of the CPF 3’-end processing/termination complex has also been implicated in termination of transcription [11,31]. Our published results demonstrated that Ssu72 exhibits a genetic and functional interaction with Rpb4, and we further showed that Ssu72 association with the chromatin and RNAPII depended on Rpb4 [29]. Since Rpb4 is functionally related to Ssu72, which is a critical gene looping and 3’-end processing/termination factor, we wondered if Rpb4 also participates in gene loop formation and termination of transcription in budding yeast.

We therefore examined the conformation of three constitutively expressed genes *HEM3*, *BLM10* and *SEN1* in *rpb4Δ* and isogenic wild type (*wt*) cells. Gene looping was detected by the ‘Chromosome Conformation Capture’ (3C) approach. This procedure converts the promoter-terminator interaction into quantitatively measurable PCR products obtained using the primer pair flanking the promoter and terminator region, as schematically represented in **Figure 1A**. 3C analysis was performed following the protocol described by [32]. As expected, the 3C assay detected a strong looping interaction due to physical association of the promoter and terminator regions of *BLM10*, *HEM3* and *SEN1* in *wt* cells (**Figure 1B**, black bars). In the absence of Rpb4, however, looping interaction decreased by about 40-70% (**Figure 1B**, grey bars). As expected, transcription of these genes, as determined using reverse transcription-PCR (RT-PCR), exhibited a declining trend in *rpb4Δ* cells relative to *wt* cells (**Figure 1C**). We have previously examined the effect of the *rpb4Δ* mutation on two inducible genes, *INO1* and *MET16*. In the absence of Rpb4, transcription of *INO1* decreased while that of *MET16* remained unaffected [33]. 3C analysis exhibited a similar effect of *rpb4Δ* mutation on gene looping. Formation of gene loops, as determined in terms of P1T1 PCR product, was compromised for *INO1* but not *MET16* in *rpb4Δ* cells compared to the isogenic *wt* cells [33]. On the basis of these results, it is reasonable to conclude that Rpb4 may not be a general gene looping factor like TFIIB, but is affecting the transcription and gene looping of a subset of RNAPII-transcribed genes.

**FIGURE 1.**
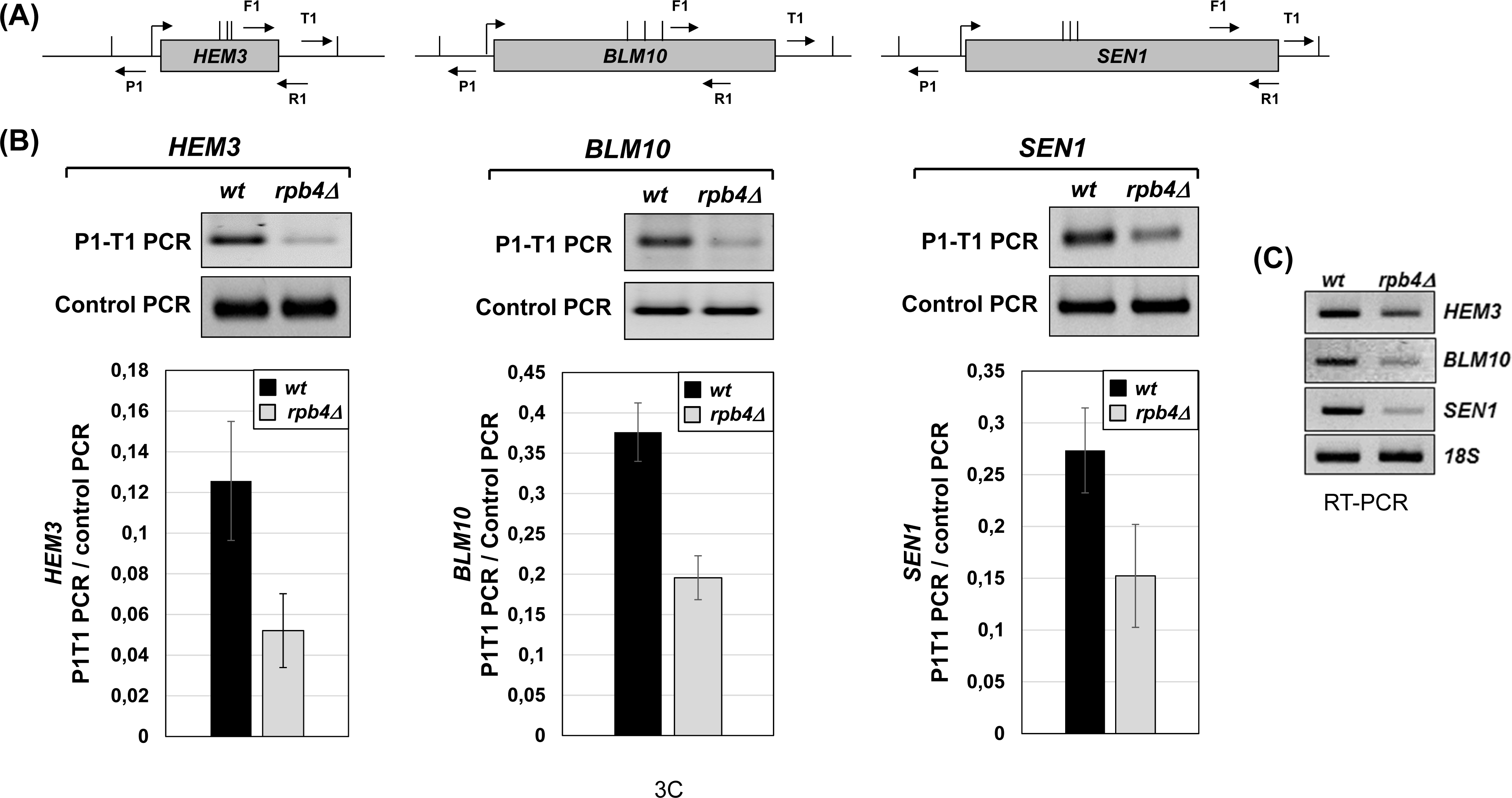
3C analysis of *HEM3*, *BLM10*, and *SEN1* genes in *rpb4Δ* and isogenic wild type cells. (A) Schematic representation of *HEM3*, *BLM10*, and *SEN1* genes indicating the positions of P1 and T1 primers used for 3C analysis and F1R1 primers used for control PCR. (B) 3C results for *HEM3*, *BLM10*, and *SEN1* genes in *rpb4Δ* and wild type cells (*wt*). P1T1 PCR reflects the looping signal and the F1R1 PCR reflects the loading control indicating that an equal amount of template DNA was used in each 3C PCR reactions. Lower panel showing the quantification of results shown above in gels, has P1T1 PCR signals normalized by control PCR. Quantifications represent the results of at least three biological replicates. Error bars indicate one unit of standard deviation. (C) RT-PCR analysis of *HEM3*, *BLM10*, and *SEN1* in *rpb4Δ* and wild type cells.

Since gene looping occurs in a transcription-dependent manner [17], it can be presumed that all mutations that negatively affect transcription would in principle affect gene looping. If this logic is true, then RNAPII will be playing a rather passive role in gene loop formation, and the observed gene looping defect in *rpb4Δ* cells could be an indirect consequence of the transcription defect. Our data, however, demonstrate that this is not the case as not all RNAPII mutants exhibit a gene looping defect. 3C analysis revealed that *rpo21-4* [34,35] and *rpb2-4* [36], which are mutants of *RPB1* and *RPB2* subunits of RNAPII respectively, do not exhibit a measurable gene looping defect (**Figure EV1C-B**). These results were corroborated by our previous observation that mutations in Rpb3 *(rpb3-E6K)* and Rpb11 (*rpb11-E108G*) subunits of RNAPII also had no effect on gene looping [33]. A logical interpretation of these data is that Rpb4, unlike other RNAPII subunits, may be an active player in gene loop formation in budding yeast.

**FIGURE EV1.**
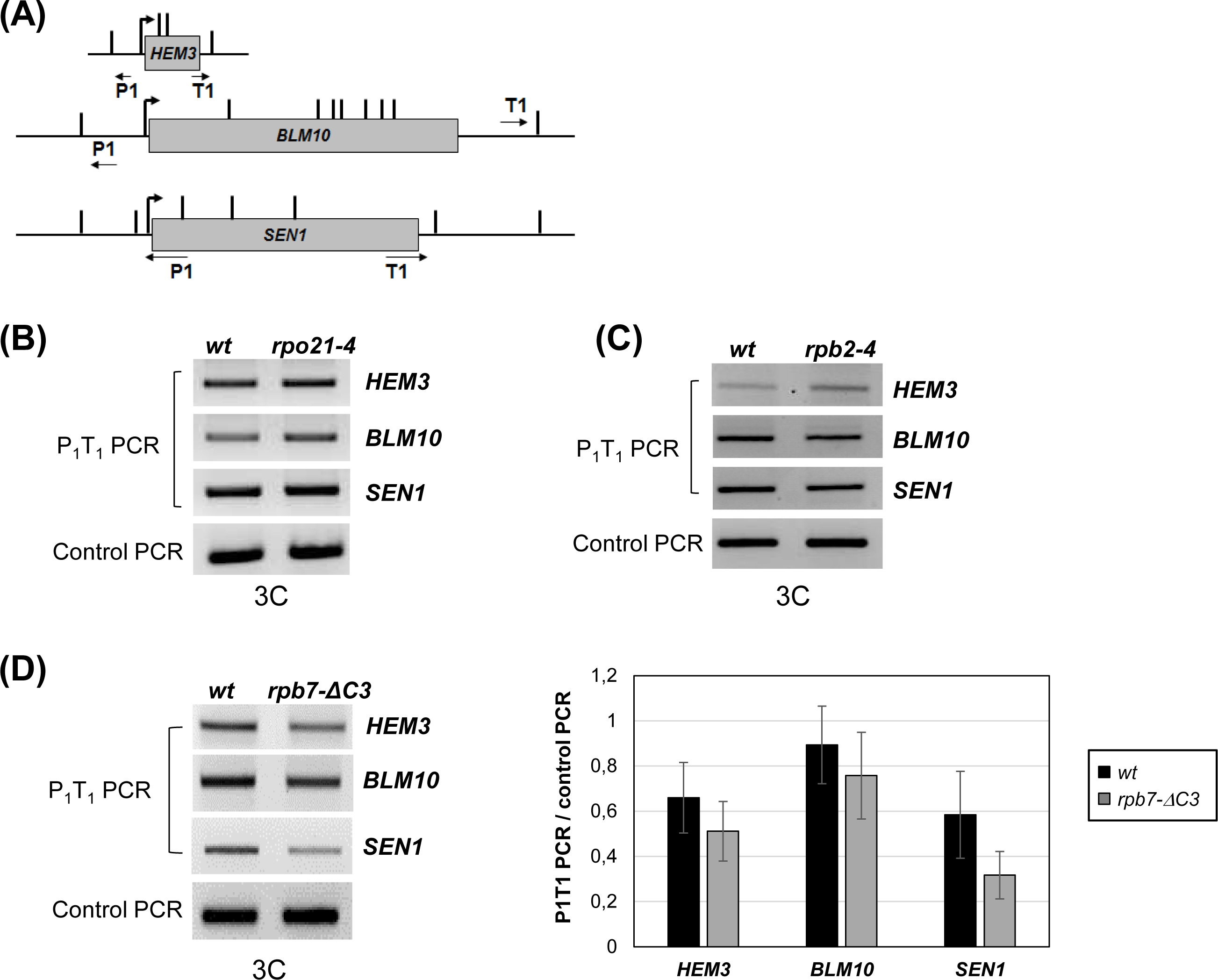
3C analysis of *HEM3*, *BLM10*, and *SEN1* genes in several RNAPII subunit mutants; *rpo21-4*, *rpb2-4* and *rpb7-ΔC3*. (A) Schematic representation of *HEM3*, *BLM10*, and *SEN1* genes indicating the positions of P1 and T1 primers used for 3C analysis. (B) 3C results for *HEM3*, *BLM10*, and *SEN1* genes in *rpo21-4* and wild type (*wt*) cells. P1T1 PCR reflects the looping signal and the control PCR represents an intergenic region of chromosome V used, used as loading control. PCR products were run in a 1.5% agarose gel stained with EtBr. (C) Same as in (B) for *rpb2-4* and wild type (*wt*) cells. (D) Left panel; 3C results for *HEM3*, *BLM10*, and *SEN1* genes in *rpb7-ΔC3* and wild type (*wt*) cells. Right panel; quantification of 3C results, where P1T1 PCR signals were normalized by control PCR signals. Error bars represent standard deviations.

To probe the issue further, we performed 3C analysis in *rpb7Δ3*, which is a mutant of the *RPB7* subunit of RNAPII. Rpb7 along with Rpb4 forms the stalk of RNAPII and the *rpb7Δ3* mutation, in addition to reduced transcription, has an adverse effect on association of Rpb4 with transcriptionally active chromatin [29]. Our results show that there is only a marginal decrease in looping PCR signal in *rpb7Δ3* cells compared to the isogenic wild type strain (**Figure EV1D**). Although association of Rpb4 with the chromatin is compromised in *rpb7Δ3*, even this reduced amount of chromatin-linked Rpb4 would be sufficient to confer looped gene conformation. A similar explanation holds true for *rpo21-4*, which is a mutant in the Rpb1 subunit located in the foot domain of RNAPII and affects the stability as well as the assembly of the RNAPII complex. In fact, in the *rpo21-4* mutant, assembly of Rpb4/7 into RNAPII and transcription levels are decreased [34].

Our data strongly suggest that Rpb4 regulates the transcription of a subset of genes through its direct role in gene loop formation. The question now is to elucidate the factors that interact with Rpb4 and allow it to perform its function in the regulation of transcription.

### TFIIB association to the RNAPII complex is impaired in the absence of Rpb4

The general transcription factor TFIIB is a key player in the formation of gene loops [17]. TFIIB is an evolutionarily conserved transcription factor that plays an essential role in initiation of transcription. Recent studies, however, have implicated TFIIB in the termination step of transcription as well. TFIIB involvement in termination has been observed in yeast [12,37], mammals [7] and flies [6]. In yeast, TFIIB is encoded by the *SUA7* gene and, specifically, the *sua7-1* mutation negatively affects transcription start site selection [38], transcription termination [12] and severely impairs formation of genes loops [10,17]. We carried out 3C analysis to compare the extent of loss of gene looping in *rpb4Δ* and *sua7-1* mutants. We observed that loss of gene looping measured in terms of P1T1 PCR product was comparable in *rpb4Δ* and *sua7-1* cells (**Figure 2**, and [17]. Both Rpb4 and TFIIB also exhibit similarity in terms of promoter occupancy and genetic interaction with 3’end processing-termination factors [29,37,39]. We next examined if Rpb4 influences the association of TFIIB with RNAPII.

**FIGURE 2.**
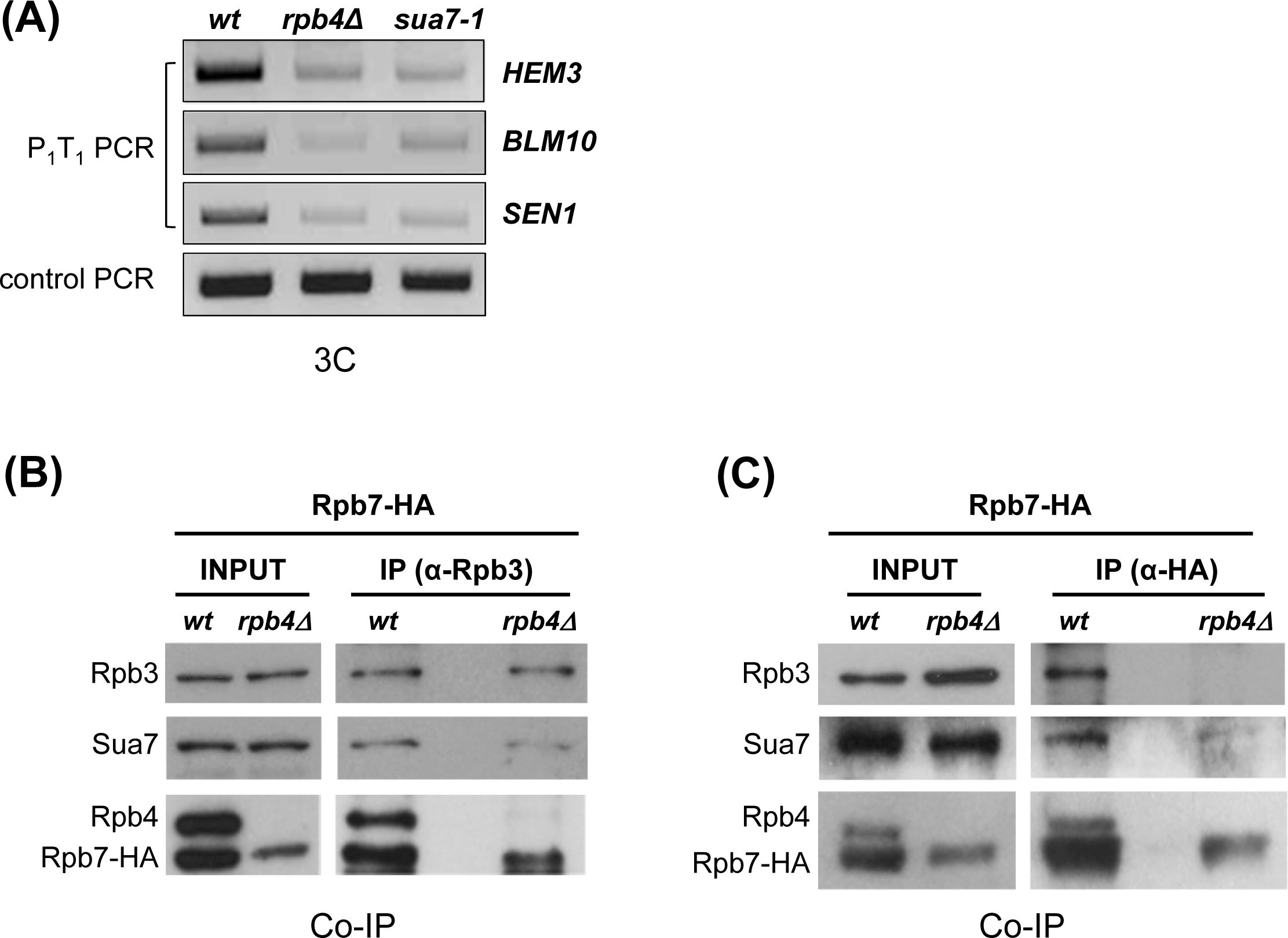
TFIIB association to the RNAPII complex is impaired in the absence of Rpb4. (A) 3C data showing that *rpb4Δ* and *sua7-1* cells exhibit loss of gene looping to a similar extent when compared to *wt* cells. PCR was performed with P1T1 divergent primers for *HEM3*, *BLM10* and *SEN1* genes. Control PCR represents an intergenic region of chromosome V was used as loading control. PCR products were run on a 1.5% agarose gel stained with EtBr. (B) TFIIB (Sua7) association with RNAPII is significantly reduced in *rpb4Δ* cells when the polymerase is immunoprecipitated with anti-Rpb3. Co-immunoprecipitations (Co-IP) were performed with WCEs from *wt* and *rpb4*Δ strains expressing Rpb7-HA using anti-Rpb3. Input and IPs were analyzed by Western blotting with antibodies to the indicated proteins. (C) TFIIB (Sua7) and Rpb3 association with Rpb7-HA is extraordinarily reduced in *rpb4Δ* cells. The Co-IPs were performed with WCEs from *wt* and *rpb4*Δ strains expressing Rpb7-HA using anti-HA. Input and IPs were analyzed as in (B).

The crystal structure of the RNAPII-TFIIB complex does not suggest a direct role of Rpb4 in TFIIB-RNAPII interaction during assembly of the preinitiation complex (PIC) [40,41]. However, the 12-subunit RNAPII complex may be a requisite for the stable association of TFIIB with a termination-competent RNAPII. Since terminator occupancy of TFIIB is dependent on Ssu72 [17], and Ssu72 association with the chromatin, in turn, depends on Rpb4 [29], we decided to investigate whether Rpb4 is required for an optimal TFIIB-RNAPII interaction. We immunoprecipitated RNAPII using antibodies against Rpb3 from whole cell extract of *wt* and *rpb4Δ* cells and detected the presence of TFIIB (Sua7) in the coimmunoprecipitated fraction by Western blot (**Figure 2B**). Coimmunoprecipitation was performed in strains expressing Rpb7-HA, so that Rpb7 and Rpb4 levels can be tested as controls in the experiment. As shown in **Figure 2B**, almost equal levels of Rpb3 were immunoprecipitated from *wt* and *rpb4Δ* extracts, but a significant reduction in TFIIB (Sua7) level associated with RNAPII was observed in the absence of Rpb4. As reported earlier, Rpb7-HA levels also decreased in *rpb4Δ* cells [42]. We next pulled down Rpb7 from a strain harboring HA-tagged Rpb7 using anti-HA antibody and looked for the presence of TFIIB and Rpb3 in the immunoprecipitated fraction. As expected, we detected TFIIB, Rpb3 and Rpb4 coimmunoprecipitating with Rpb7 in *wt* cells (**Figure 2C**). In the absence of Rpb4, however, there was a dramatic decline in the amount of TFIIB associated with Rpb7 while Rpb3, representing the 10-subunit RNAPII, was undetectable (**Figure 2C**). This is consistent with the fact that Rpb4 is required for Rpb7 association with the RNAPII complex [19,39,43]. In addition, these data support the idea that a complete 12-subunit RNAPII complex could be required for the stable association of TFIIB with a competitive RNAPII termination complex.

### Rpb4 does not directly affect termination of transcription

All the termination-defective mutants of CF1A and CPF 3’-end processing complexes are defective in gene looping [37,44]. The mutants of general transcription factors TFIIB, TFIIH and Mediator complex, which have only a moderate effect on initiation but exhibit a terminator readthrough phenotype, are also defective in gene looping [12,16]. This strong correlation between gene looping and termination of transcription prompted us to investigate if Rpb4, owing to its involvement in gene looping, is playing a role in termination of transcription. Therefore, we performed strand-specific TRO analysis for *BLM10*, *HEM3* and *SEN1* in *rpb4Δ* and isogenic *wt* cells. We expected that if Rpb4 is required for termination of transcription, a terminator-readthrough phenotype will be observed in *rpb4Δ* cells. Our results show that there was no appreciable TRO signal beyond the poly(A) signal in *wt* cells for all three genes (region 2 in **Figure 3B**, black bars). An almost identical TRO signal pattern was observed in *rpb4Δ* cells (**Figure 3B**, white bars). These results strongly suggest that Rpb4, despite playing an active role in gene looping, is not directly involved in termination of transcription of at least three genes used in this investigation.

**FIGURE 3.**
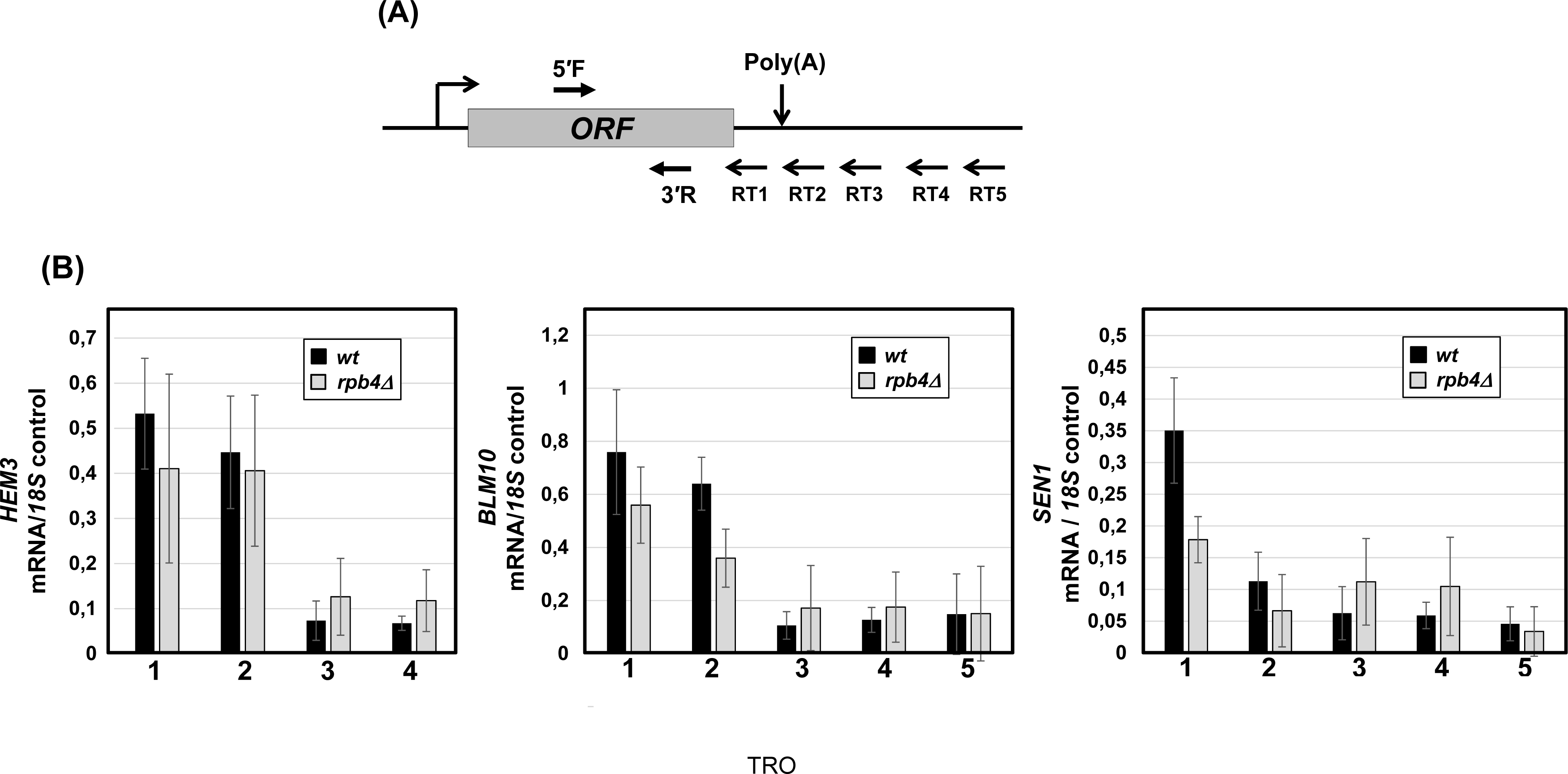
Transcription Run-On (TRO) analysis of *HEM3*, *BLM10*, and *SEN1* genes in *rpb4Δ* and isogenic wild type cells. (A) Schematic representation of *HEM3*, *BLM10*, and *SEN1* genes indicating the positions of RT1, RT2, RT3, RT4 and RT5 primers used for cDNA synthesis, as well as 5’F and 3’R primers used for PCR amplification of cDNA. (B) Quantification of RNA levels detected following TRO analysis in *rpb4Δ* and wild type (*wt*) cells. The transcript level of *18S* was used as the control for normalization of results. Quantifications represent the results of four biological replicates. Error bars indicate one unit of standard deviation.

### Rpb4 overexpression overcomes the gene looping and termination defect of *sua7-1*

Our results clearly demonstrate that Rpb4 plays a critical role in facilitating interaction of TFIIB with RNAPII. This prompted us to explore if Rpb4 has any bearing on the transcription-related functions of TFIIB in yeast cells, such as gene looping and termination of transcription [12,17]. Therefore, we first examined if Rpb4 affects TFIIB role in gene looping of *HEM3*, *BML10* and *SEN1.* We transformed the *sua7-1* mutant with a plasmid overexpressing Rpb4 and a control plasmid, and monitored looped gene architecture of *HEM3*, *BML10* and *SEN1* by 3C approach. Our results show that the looped conformation of all three genes, measured in terms of P1T1 PCR product, was as expected severely compromised in *sua7-1* cells **(Figure 4B and 4C**, grey bars). Upon overexpression of Rpb4, however, gene looping of *HEM3* and *BML10* was completely restored to wild type level, while that of *SEN1* was partially restored (**Figure 4B and 4C**, white bars). These results clearly demonstrate that *RPB4* overexpression is able to reverse the gene looping defect of the TFIIB mutant, and strongly support a functional connection between Rpb4 and TFIIB during the formation of gene loops.

**FIGURE 4.**
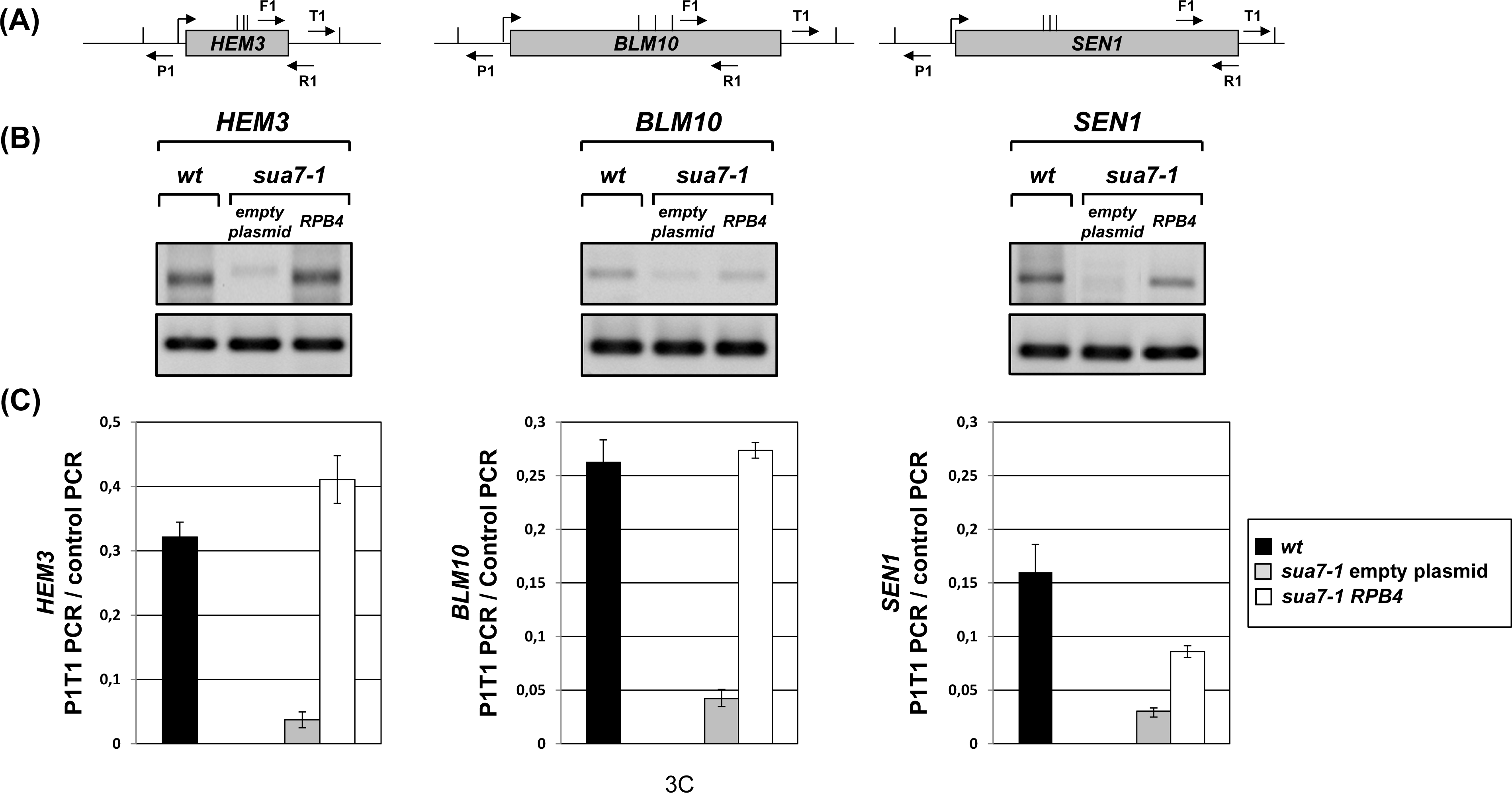
3C analysis of *HEM3*, *BLM10*, and *SEN1* genes in wild type, *sua7-1* and *sua7-1* cells overexpressing *RPB4*. (A) Schematic depiction of three genes used in this analysis indicating the positions of P1 and T1 primers used for 3C PCR and F1-R1 primers used for control PCR. (B) Gels showing 3C PCR results for *HEM3*, *BLM10*, and *SEN1* in wild type cells and *sua7-1* cells without overexpression of Rpb4 (empty plasmid) or with overexpression of Rpb4 (pRPB4). P1T1 PCR reflects gene looping and the F1R1 PCR reflects the loading control indicating that an equal amount of template DNA was used in each 3C reaction. (C) Quantification of results shown above in gels has P1T1 PCR signals normalized with respect to the control PCR. Quantifications represent the results of at least three biological replicates. Error bars indicate one unit of standard deviation.

We next investigated if Rpb4 has any influence on TFIIB-mediated termination of transcription, though it seems that deletion of *RPB4* has no negative effect on this process (**Figure 3**). We used strand-specific TRO approach to monitor termination of transcription of *HEM3*, *BML10* and *SEN1* in wild type cells, *sua7-1* mutant and *sua7-1* mutant overexpressing Rpb4 as described above. In wild type cells, there was no detectable TRO signal beyond the poly(A) site for all three genes thereby demonstrating efficient termination of transcription (**Figure 5B**, white bars). In *sua7-1* cells, however, TRO signal was detected beyond the poly(A) site in the regions 2, 3 and 4 for *SEN1* as well as regions 2 and 3 for *BLM10* and *HEM3* (**Figure 5B**, black bars). Clearly, the polymerase was unable to read the termination signal in *sua7-1* cells and readthrough in the regions downstream of the poly(A) site. The termination-defective phenotype of the mutant, however, was almost completely rescued upon overexpression of Rpb4 (**Figure 5B**, grey bars). Just like in the case of gene looping described above, overexpression of *RPB4* overcame the transcription termination defect in *sua7-1* cells. Taken together, these data strongly reinforce a functional connection between Rpb4 and TFIIB during gene loop formation and termination of transcription.

**FIGURE 5.**
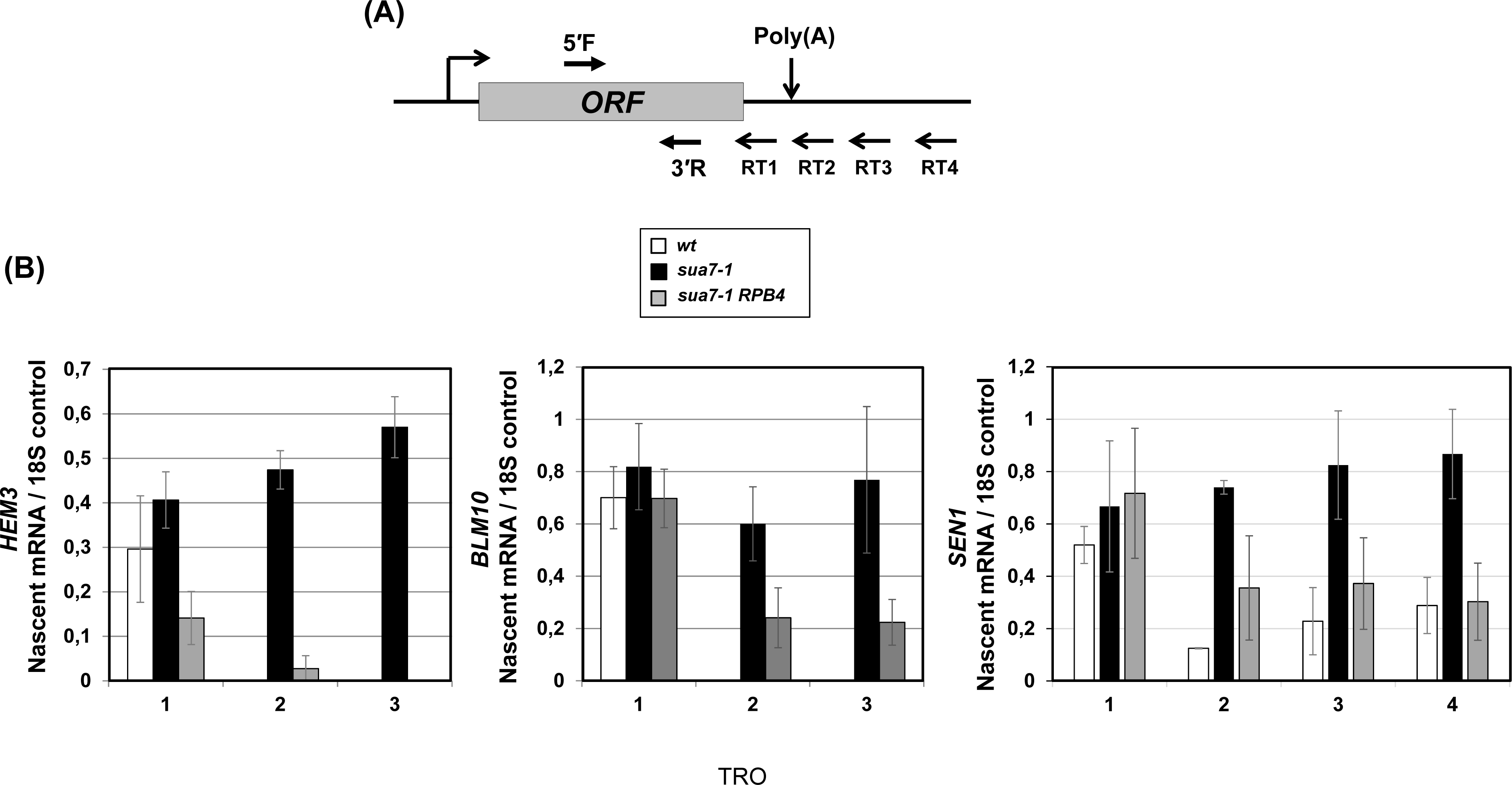
TRO analysis of *HEM3*, *BLM10*, and *SEN1* in wild type, *sua7-1* and *sua7-1* cells overexpressing *RPB4*. (A) Schematic representation of *HEM3*, *BLM10*, and *SEN1* genes indicating the positions of RT1, RT2, RT3 and RT4 primers used for cDNA synthesis, as well as 5’F and 3’R primers used for PCR amplification of cDNA. (B) Quantification of RNA levels detected following TRO in wild type cells and *sua7-1* cells without overexpression of Rpb4 (empty plasmid) or with overexpression of Rpb4 (pRPB4). The transcript level of *18S* was used as the control for normalization of results. Quantifications represent the results of at least three biological replicates and six technical replicates. Error bars indicate one unit of standard deviation.

### Rpb4 and Ssu72 are functionally linked during the formation of gene loops

Another major player involved in gene looping is Ssu72 [10,17]. We provided a genetic and functional connection of Ssu72 with Rpb4 [29]. We observed that *RPB4* overexpression was able to rescue the temperature-sensitive phenotype of *ssu72-2* quite well, and *SSU72* overexpression partially rescued the growth defect of *rpb4* mutant at 28 °C and 33°C (**Figure 2EVA-B**; and [29]). Furthermore, overexpression of *SSU72* suppressed the effect on CTD dephosphorylation observed in the *RPB4* deletion mutant [29]. Then, we decided to test by 3C whether the suppression of growth defect was due to restoration of gene looping. We, therefore, tested if *SSU72* overexpression can overcome the gene looping defect of *rpb4Δ* cells. We transformed *rpb4Δ* and isogenic wild type strains with a high copy plasmid overexpressing *SSU72*. Our results clearly show that increased Ssu72 levels effectively restored the impaired loop conformation in *HEM3*, *BLM10* and *SEN1* genes, measured in terms of P1T1 PCR product, in *rpb4Δ* cells (**Figure 6A**, white bars). This is consistent with the suppression of *rpb4Δ* growth defects upon overexpression of *SSU72* (**Figure S2A**, and [29]). We next performed 3C assays in *wt* and *ssu72-2* cells overexpressing *RPB4*. The *ssu72-2* strain displays thermosensitivity at 37°C (**Figure EV2B**, and [29,30] and has been shown to exhibit a gene looping defect [30]. Our results show that although in *ssu72-2* cells the gene looping defect is less pronounced than in *sua7-1* or *rpb4Δ* cells (**Figure 2A**), the overexpression of *RPB4* (**Figure 6B**), or of *RPB4* and *RPB7* together (**Figure EV2C**), completely restored the looped conformation. These results are consistent with the suppression of the thermosensitivity of *ssu72-2* at 37°C by increasing *RPB4* levels (**Figure EV2B**).

**FIGURE EV2.**
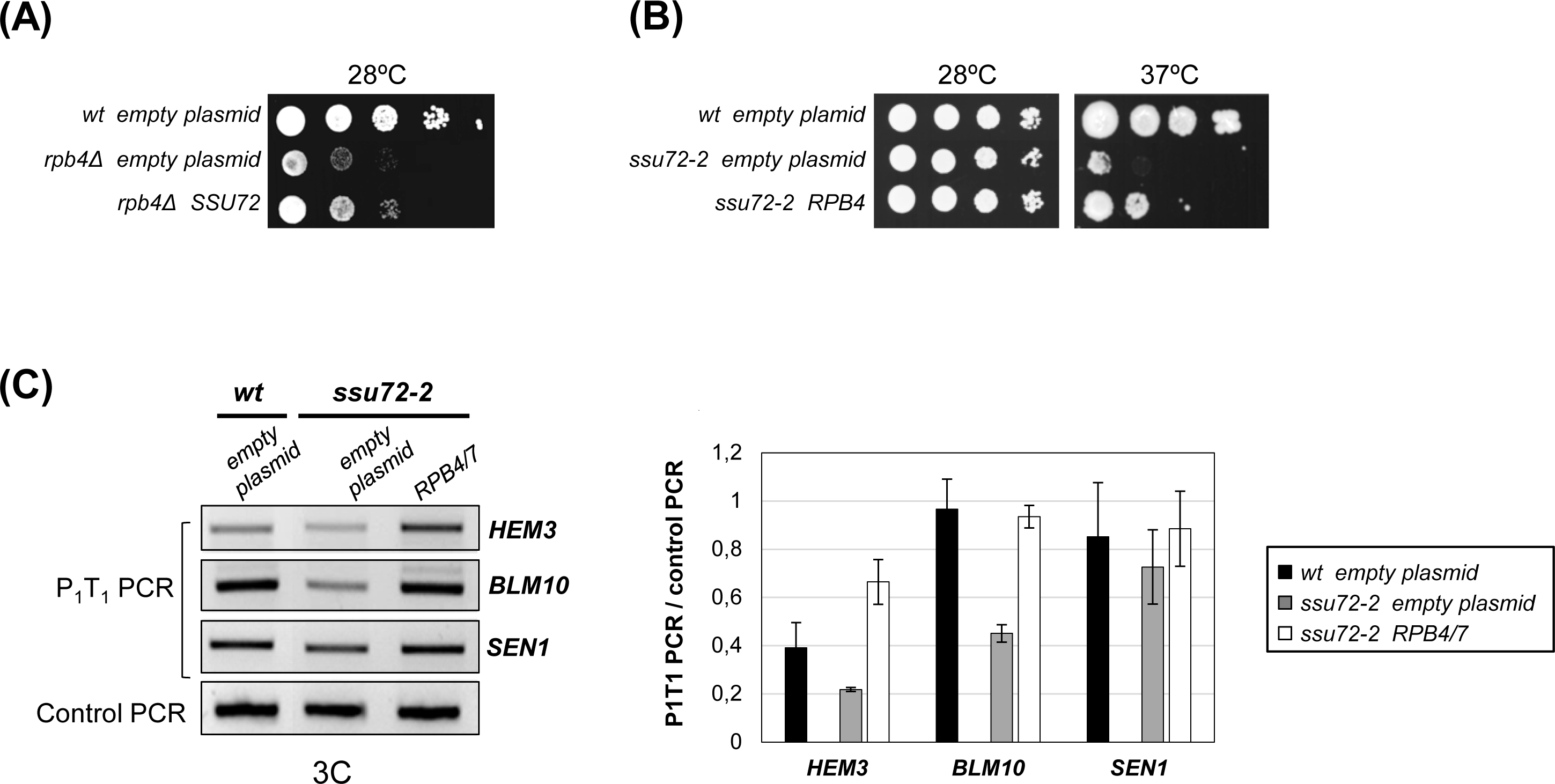
Genetic and functional interaction between *RPB4* and *SSU72*. (A) Overexpression of *SSU72 (pSSU72)* suppresses the slow growth of the *rpb4Δ* strain at 28°C, as previously reported [29]. *wt* strain was transformed with an empty plasmid and *rpb4Δ* strains was transformed with an empty plasmid or with a high-copy plasmid containing *SSU72.* Serial dilutions (1:10) of the strains tested were spotted on SC medium and grown for 2-3 days at 28°C. (B) Overexpression of *RPB4* suppress the thermo-sensitivity of the *ssu72-2* strain. The *wt* strain was transformed with an empty plasmid and *ssu72-2* was transformed with an empty plasmid or with a high-copy plasmid bearing *RPB4.* Serial dilutions (1:10) of the indicated strains were spotted on SC media for 2-3 days at 28 and 37°C. (C) Overexpression of *RPB4* and *RPB7* (*RPB4/7*) in *ssu72-2* suppresses gene looping defects. The wild-type (*wt*) strain was transformed with an empty plasmid and *ssu72-2* strain was transformed either with an empty plasmid or with high-copy plasmids bearing *RPB4* or *RPB7*. Gene looping was assayed by 3C and PCR products were run in a 1.5% agarose gel stained with EtBr. P1T1 PCR products were quantified by dividing P1T1 PCR signals by control PCR signals for each sample and graphed. Error bars represent standard deviation.

**FIGURE 6.**
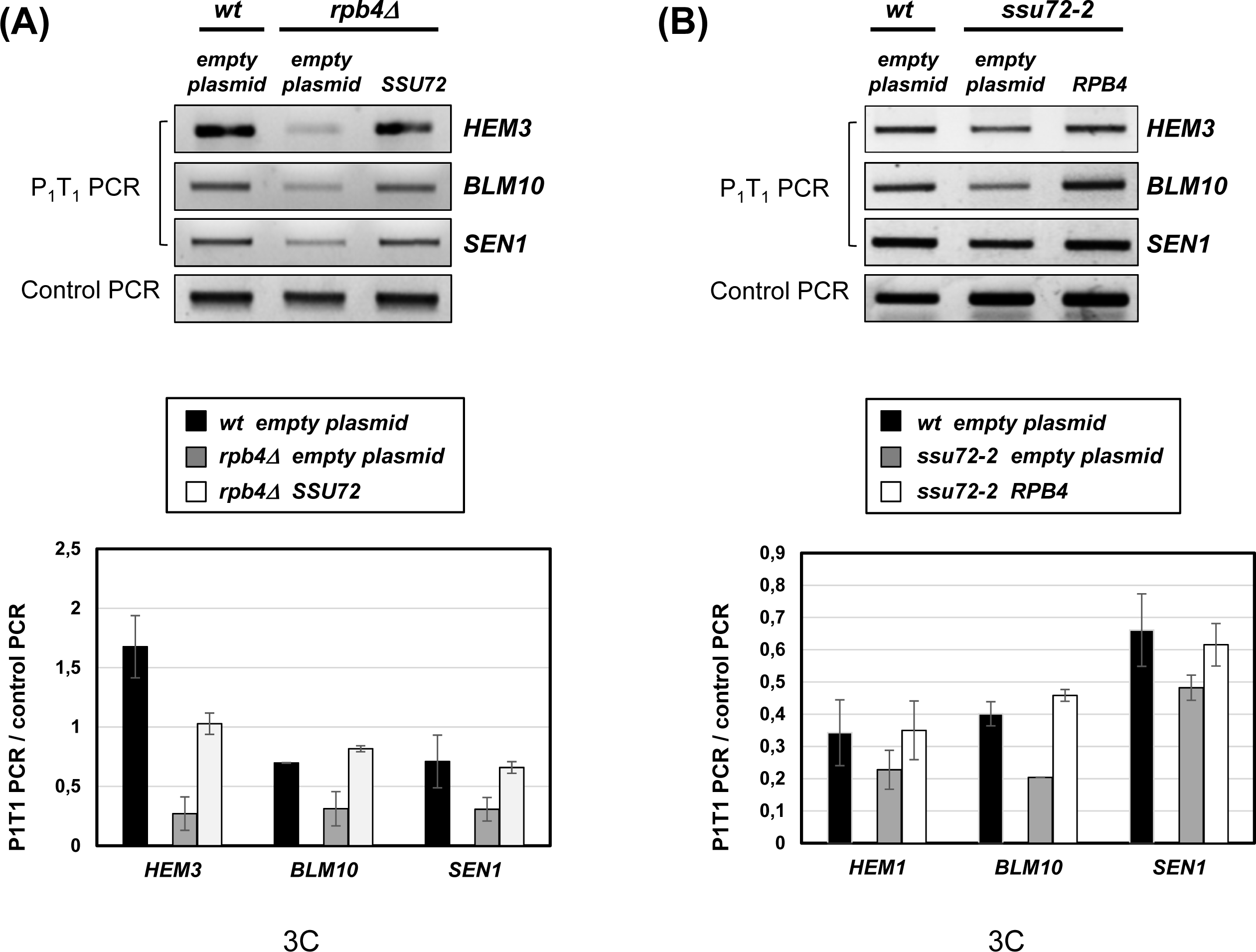
Rpb4 and Ssu72 are functionally linked during the formation of gene loops. (A) Overexpression of *SSU72* in *rpb4Δ* cells restores the formation of gene loops to *wt* levels. The *wt* strain was transformed with an empty plasmid and *rpb4Δ* strain was transformed either with an empty plasmid or with a high copy plasmid expressing *SSU72*. Gene looping was assayed by 3C and PCR products were run in a 1.5% agarose gel stained with EtBr. P1T1 PCR products were quantified by dividing P1T1 PCR signals by control PCR signals for each sample. Graph in the lower panel represent the mean values from the quantification of three experiments, where error bars represent standard deviations. (B) *RPB4* overexpression in *ssu72-2* cells restores the formation of gene loops to *wt* levels. The *wt* strain was transformed with an empty plasmid and *ssu72-2* strain was transformed either with an empty plasmid or with a high copy plasmid expressing *RPB4*. Quantifications and representations where performed as in (A).

The interaction of Ssu72 with the Ser5P and Ser7P residues of the Rpb1-CTD, which are the targets of its phosphatase activity, is well established [8,9,45]. A physical interaction of Ssu72 with the Rpb2 subunit has also been reported [46]. The data presented here and in our published report [29], supports a direct interaction between Rpb4 and Ssu72. We, therefore, wondered whether Rpb4 (or the Rpb4/7 heterodimer) could be the direct target of Ssu72 within the 12-subunit RNAPII molecule, and whether this interaction promotes loop conformation. To check our hypothesis, we expressed and purified GST-Ssu72 and Rpb4/Rpb7-6His recombinant proteins from *E. coli* and performed *in vitro* interaction experiments. We could not detect a direct physical interaction between the CTD phosphatase and the heterodimer (data not shown). This could be either due to absence of any direct interaction between Ssu72 and Rpb4 or that the interaction between these two proteins depends on something else not present in the reaction mix. We then decided to use another strategy to answer the question, and immunoprecipitated the Rpb4/Rpb7 heterodimer from whole cell extracts of a strain expressing Rpb7-HA using anti-HA antibody. Afterwards, we incubated the immunoprecipitated proteins with either GST or GST-Ssu72 (**Figure 7A**). Our results show that Rpb4/7 specifically interact with Ssu72, and that this interaction is lost in the absence of Rpb4. In addition, we submitted the immunoprecipitated proteins for mass spectrometric analysis. Ssu72 and TFIIB were identified among the proteins interacting with Rpb7-HA (data not shown). These data strongly indicate that Rpb4 and Ssu72 may be the part of the same complex during the formation of gene loops.

**FIGURE 7.**
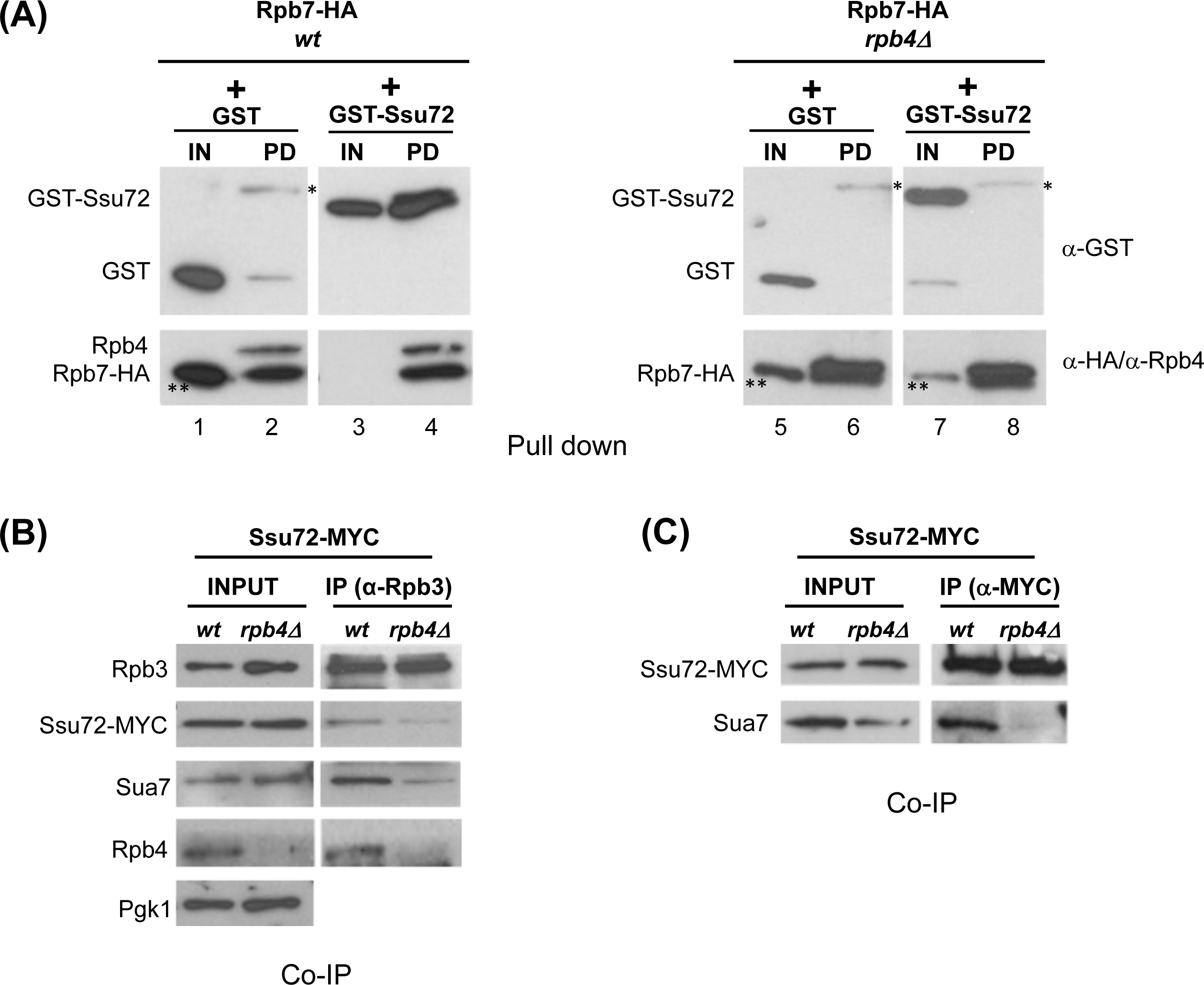
TFIIB association with RNAPII and to Ssu72 is impaired in the absence of Rpb4. (A) Pull down assay showing the interaction between Ssu72 and the Rpb4/7 heterodimer. Rpb7-HA was immunoprecipitated from *wt* and *rpb4Δ* whole cell extracts (WCE) using the anti-HA antibody. Double amount of WCE were used in the case of *rpb4Δ*, because Rpb7-HA levels are lower in that mutant (See figure 2B; [42]). Thereafter, the immunoprecipitated proteins were incubated either with GST or GST-Ssu72 recombinant proteins. After extensive washing, the interaction of Rpb7-HA with the recombinant proteins (PD) was assayed by Western blotting with the corresponding antibodies. In the input (IN) lanes (1, 3, 5 and 7) the same amount of GST or GST-Ssu72 used in the pull down assays were loaded. (*) unspecific band. (**) Signal remaining from anti-GST blotting. The GST protein has similar size to Rpb7-HA (around 25 KDa). The membranes were first incubated with anti-GST, and then washed and incubated with a mix of anti-HA and anti-Rpb4 antibodies. GST-Ssu72 suffered proteolytic cleavage and we detected some GST in the input lane (lane 7) when blotting with anti-GST antibody. (B) Association of Ssu72 and TFIIB (Sua7-MYC) with RNAPII is reduced in *rpb4*Δ cells. Co-IPs performed from WCEs of Ssu72-MYC cells (*wt* and *rpb4*Δ) using anti-Rpb3 antibodies. Input and IPs were analyzed by Western blotting with antibodies to the indicated proteins. (C) The association of TFIIB (Sua7) with Ssu72 is hardly detectable in *rpb4*Δ cells. Co-IPs were performed from WCEs of Ssu72-MYC cells (*wt* and *rpb4*Δ) using anti-MYC antibody. Input and IPs were analyzed by Western blotting with antibodies to the indicated proteins.

### Rpb4 is required for the stable association of Ssu72 and TFIIB

Next, based on the functional link between Rpb4 and the two gene looping factors, Ssu72 and TFIIB, who also exhibit functional interaction with each other [11,47,48], we investigated whether Rpb4 influences the interaction between them. A positive result would explain why *rpb4Δ* cells present such a dramatic defect on gene loops formation. To address the issue, we prepared whole cell extracts from *wt* and *rpb4Δ* strains expressing Ssu72-MYC. First, we analyzed the levels of Ssu72-MYC and TFIIB proteins associated with RNAPII by immunoprecipitating Rpb3 (**Figure 7B**). In agreement with our published results [29], Ssu72-MYC association with RNAPII is reduced in the absence of Rpb4 (**Figure 7B**). This is reminiscent of the result presented in Figure 2A, where the interaction of TFIIB with the 10-subunit polymerase complex is also impaired in the cells lacking Rpb4 (**Figure 7B**). Very interestingly, when we immunoprecipitated Ssu72-MYC, though similar amounts of Ssu72-MYC were detected in both *wt* and *rpb4Δ* cells, amount of TFIIB associated with it was dramatically reduced and was barely detectable in *rpb4Δ* cells (**Figure 7C**). In other words, in the absence of Rpb4, the complex formed by TFIIB and Ssu72 is impaired. In conclusion, Rpb4 is required for the stable association of Ssu72 with TFIIB for facilitating gene loop conformation. Altogether, our data strongly indicate that Rpb4 works in concert with Ssu72 and TFIIB to promote the formation of gene loops (**Figure 8**), and that RNAPII is not simply a passive actor in this important regulation process.

**FIGURE 8.**
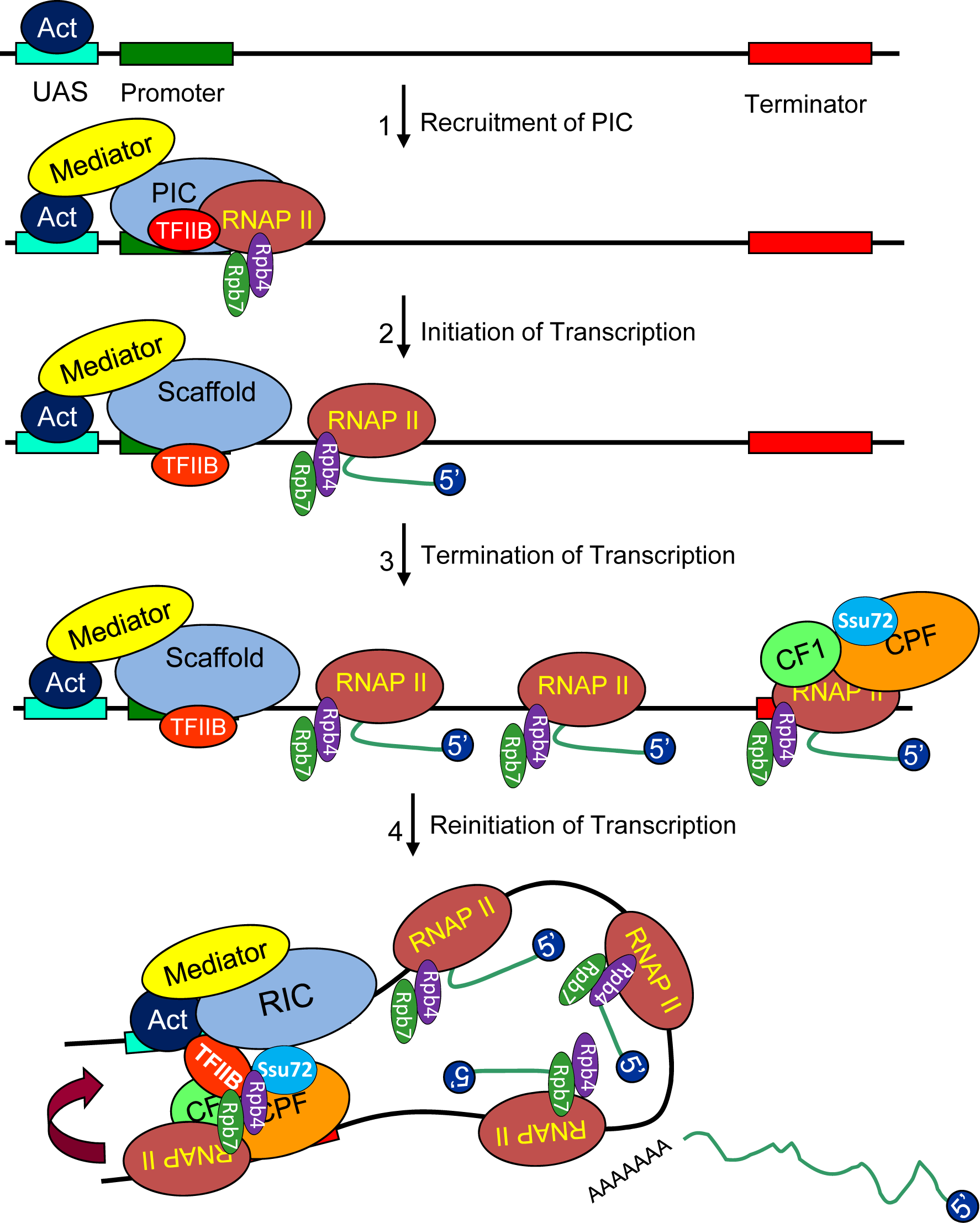
A working model for Rpb4-dependent gene looping. Rpb4 facilitates the association of Ssu72 with the promoter-bound TFIIB at the end of pioneer round of transcription. TFIIB-Ssu72 interaction results in gene loop formation. The proximity of the promoter and terminator region in the gene loop may help in transfer of polymerase from the terminator to the promoter for reinitiation of transcription. Act=activator, RIC=reinitiation complex.

## DISCUSSION

Genes can be viewed, in a simplistic manner, as a linear DNA sequence containing promoter and terminator regions at the 5’ and 3’ ends, respectively, where RNAPII initiates and terminate transcription. Upon termination of transcription, RNAPII stops and dissociates from the gene. At this point, RNAPII needs to be recycled to reinitiate a new round of transcription. Therefore, an efficient system to recycle the polymerase will contribute to maintaining faithful gene expression. In that sense, juxtaposition or direct interaction between the promoter and terminator by gene looping, facilitated by the physical association of initiation and termination factors, promotes efficient RNAPII recycling, transferring the enzyme to the promoter to reinitiate transcription [49,50]. The direct involvement of RNAPII in this process has not been reported so far, though it can be envisioned that an active role of the polymerase in the formation of gene loops will substantially contribute to efficient gene expression. Nevertheless, because gene looping is a transcription dependent process [17], it can be assumed that the role of RNAPII during the formation of gene loops is simply limited to providing the first round of transcription, required for the establishment of the loop. Furthermore, it can be deduced that all the transcription mutants would present gene looping defects, as it was shown for the thermosensitive *rpb1-1* mutant, where the formation of gene loops was completely abolished when transcription was shut off at 37°C [17]. Our data, however, show on one hand that not all the RNAPII mutants with a defect in transcription display gene looping defects (for instance *rpo21-4, rpb2-4*), and on the other hand that the *rpb4Δ* mutant has dramatic effects on the formation of genes loops, as well as on transcription, at the permissive temperature of growth. One possibility is that the transcription defects observed in *rpb4Δ* cells are the consequence of impaired initiation of transcription in the first round of transcription. However, Rpb4 seems to have no influence on the association of initiation factors TBP and TFIIH with the chromatin template, and therefore it was suggested that Rpb4 is not necessary for PIC formation *in vivo* and in *vitro* [39]. Alternatively, it is conceivable that the transcription defects are the result of the inability of *rpb4Δ* cells to form gene loops and reinitiate transcription after the first round of transcription. Actually, we showed a decrease in Rpb1 occupancy not only at the 3’end of the genes in *rpb4Δ* cells in agreement with published results [39], but at the promoters as well [29]. Moreover, we found that the relative occupancy of Rpb1 at the terminator regions of genes was higher than that at the promoters in *rpb4Δ* cells. An interpretation of these results is that in the absence of Rpb4, the release of Rpb1 from the chromatin template upon termination of transcription could be impaired, which would in turn reduce RNAPII recycling thereby leading to the reduction of Rpb1 at the promoters [29]. Our data largely support this interpretation and provide evidences for a role of Rpb4 in the gene looping process while facilitating Ssu72-TFIIB association, and consequently influencing reinitiation of transcription. Thus, in the absence of Rpb4, we detected a decrease in the amount of TFIIB interacting with RNAPII, which is required for the reinitiation of transcription [51], and also reduced association of TFIIB with Ssu72. Association of TFIIB with terminators and its function in gene looping required the function of Ssu72 [17]. Moreover, we have previously demonstrated a functional link between Rpb4 and Ssu72 in regulating Rpb1-CTD dephosphorylation [29]. Complete dephosphorylation of Rpb1-CTD by Ssu72 during termination of transcription is required to reinitiate a new round of transcription [9].

### Rpb4 and Ssu72 are coupled to regulate gene looping

Rpb4 is required for proper association of Ssu72 with RNAPII and with the chromatin [29]. Importantly, we have corroborated here that the function of Rpb4 and Ssu72 are directly coupled. First, overexpression of *RPB4* not only suppresses *ssu72-2* growth, but also its gene looping defects. Second, we reported that levels of Ser7P are extraordinarily increased in *rpb4Δ*, and to a lesser extent those of Ser5P, and that *SSU72* overexpression reduces them to *wt* levels [29]. Now we have added new evidence showing that overexpression of *SSU72* restores *rpb4Δ* gene looping as well. This fits well with several studies that proposed the role of phosphatase activity of Ssu72 in promoting the formation of gene loops [10,52]. Accordingly, Ssu72 erases Ser7P marks immediately after cleavage and polyadenylation, and renders the terminating RNAPII to a hypo-phosporylated state needed for recycling back to the promoter. Thus, inactivation of Ssu72 leads to permanent phosphorylation of Ser7. This may affect the ability of RNAPII to terminate efficiently and prevent it from being recruited to the PIC. The ultimate result will be blocking of transcription initiation and lethality [9]. Then, the extraordinary high levels of Ser7P in *rpb4Δ* cells, as a result of impaired Ssu72 function, together with a reduction in the association of Ssu72 with TFIIB, may lead to a gene looping defect and will adversely affect the reinitiation of transcription.

### Rpb4 plays an indirect role in the termination of transcription

As mentioned before, there exists a network of complex interactions between initiation and termination factors that promote the formation of gene loops. Consistent with this, some initiation factors are required for transcription termination. Conversely, some termination factors have been found to associate with the 5’ end of a gene as well as with the initiation factors, and in fact have been shown to influence transcription reinitiation [44,53-57]. Therefore, it is reasonable to propose that the transcription defective mutants, that exhibit alteration in reinitiaiton or termination, exhibit a higher propensity for the gene looping defect. In fact, transcription termination is tightly coupled to gene loop formation, and all the termination mutants analyzed so far are gene looping defective as well [37,44]. Moreover, some 3’end processing/termination mutants present reduced transcription reinitiation [56]. However, *rpb4Δ* cells do not display termination defects, which seem to be a general feature of gene looping mutants, though Rpb4 contributes to cotranscriptional recruitment of 3’-end processing factors (CFIA) [39]. Actually, all the termination-defective mutants of CF1A are gene looping defective [37,44]. Nevertheless, overexpression of *RPB4* is able to rescue not only the defect on the formation of gene loops in *sua7-1* mutant, but also its termination-defective phenotype [12,17]. These results strongly suggest that Rpb4 influences termination of transcription through TFIIB. Thus, a reduction in the association of TFIIB would in consequence cause a decrease in the association of Rna14 and Rna15, an effect observed in *rpb4Δ* cells [39]. This is in agreement with a physical interaction between TFIIB and CFI, and the role of both factors at the 3’end regions of the genes for the formation of gene loops [10,17,37,44]. Indeed, the existence of a holo-TFIIB complex required for gene looping has been demonstrated. This complex contains TFIIB, CFI and the polyA polymerase Pap1, but no other GTF. The holo-TFIIB complex is not observed in *sua7-1* cells [37]. Our data altogether strongly support a functional interaction between Rpb4 and TFIIB during gene loop formation and also in termination of transcription.

### A working model: Rpb4 mediates the association between Ssu72 and TFIIB during the formation of gene loops

Ssu72-TFIIB association is crucial for the formation of gene loops [17]. TFIIB interacts with Ssu72 through its cyclin-like domain [48,58] and associates with the 3’-end/terminator region of the gene in a Ssu72 dependent manner [17]. TFIIB interacts with RNAPII, though it doesn’t make contact with Rpb4, as revealed by the RNAPII-TFIIB structure [59]. Ssu72 also interacts with RNAPII; here we show that it interacts with a recombinant Rpb4/7 heterodimer, likely through Rpb4, while previously it was reported to interact with Rpb2 [11,36] and with the CTD of Rpb1 [60]. Importantly, the data presented here clearly demonstrate that when Rpb4 is not present in the RNAPII complex, Ssu72 and TFIIB association with the polymerase is compromised and consequently TFIIB-Ssu72 interaction is adversely effected [17,48,58]. Based on the data presented here, and in previous studies [11,36,48,58,59], we propose a working model where Rpb4 has a direct role in the formation of gene loops after the first round of transcription, facilitating the stable association of Ssu72 and TFIIB with RNAPII and thus promoting the recycling of the polymerase back to the promoter from the terminator for reinitiation of transcription (see Figure 8 legend for details). Therefore, in the absence of Rpb4, the decreased association of TFIIB with Ssu72 will impair gene looping and the recruitment of the CF1 complex at the 3’-end of genes, thereby affecting transcription termination [17,37,44]. In summary, our data indicate that Rpb4 is required for the formation of a functional complex, key for the establishment of gene loops, and show that RNAPII is not simply a passive actor in this important regulatory process.

## MATERIAL AND METHODS

### Yeast strains, media and plasmids

The strains used are inventoried in Appendix Table S1. Strain construction and other genetic manipulations were performed following standard procedures [51]. Plasmids are detailed in Appendix Table S2 and primer sequences in Appendix Table S3.

### Co-immunoprecipitation and western blot analysis

Cells expressing Ssu72-MYC and Rpb4/Rpb7-HA were grown in 200 ml of rich medium to an OD_600_ of 1.0-1.2, harvested, washed with water, and then suspended in 2.0 ml of lysis buffer (20 mM HEPES pH 7.6, 200 mM KCH_3_COO^−^, 1 mM EDTA pH 8.0, glycerol 10 %) containing protease and phosphatase inhibitors. The cell suspension was flash frozen in liquid nitrogen, and then ground in a chilled mortar to a fine powder. Subsequently, the cell lysate was thawed slowly on ice and transferred to pre-chilled tubes and centrifuged at 13,200 rpm for 20 minutes. The supernatant was collected and the concentration of total proteins was assessed by measuring absorbance at 280 nm in a Nanodrop. In experiments where Rpb3, Ssu72-MYC or Rpb7-HA were immunoprecipitated, cell extracts were incubated with 5 μl of anti-Rpb3, 1-2 μl of anti-MYC or anti-HA antibodies for 3-4 hours at 4°C, previously bound to 30 μl of Protein A Sepharose CL-4B (GE Healthcare). The IPs were extensively washed with lysis buffer and beads were suspended in SDS-PAGE sample buffer. Western blot analysis was performed using the appropriate antibodies in each case: anti-Pgk1 (Pgk1, 459250; Invitrogen), anti-HA (12CA5, Roche); anti-MYC (9E10, Roche); anti-Rpb4 (2Y14; BioLegend); anti-Rpb3 (1Y26; BioLegend); anti-Sua7 (ab63909, Abcam). The ECL reagents were used for detection.

### Expression and purification of recombinant proteins

#### Recombinant Ssu72

GST-Ssu72 was expressed from the pM620 plasmid (a gift of Dr. Hampsey) in *E. coli* BL21 (DE3) for 15h at 18°C. Subsequently, cells were harvested by centrifugation, resuspended in binding buffer (20 mM Tris pH 7.3, 150 mM NaCl, 1 mM DTT), containing EDTA-free Protease Inhibitor Cocktail (Complete™, Roche) and DNase I (Roche). Cells were sonicated, centrifuged and the supernatant loaded into a GSTrap FF (GE Healthcare) chromatography column, previously equilibrated with binding buffer. The protein was eluted with elution buffer (20 mM Tris pH 7, 150 mM NaCl, 5 mM DTT, 10 mM reduced glutathione, pH 8) in 14 column volumes and collected fractions were analyzed by SDS-PAGE. Selected fractions were pooled, concentrated in a 30 kDa cut-off centrifugal filter (Millipore) and loaded onto a MonoQ column equilibrated in start buffer (20 mM Tris pH 7.4, 100 mM NaCl, 1mM DTT). Afterwards, the protein was eluted with elution buffer (20 mM Tris pH 7.4, 1M NaCl, 1mM DTT). The fractions containing the eluted protein were collected and analyzed by SDS-PAGE. The concentrated pool of the fractions containing the protein was frozen in liquid nitrogen and stored at −80°C.

#### Pull down-assays

Cells were grown to an OD600 of 0.8, collected, washed and suspended in lysis buffer (20 mM HEPES pH 7.6, 200 mM KCH_3_COO^−^, 1 mM EDTA pH 8.0, glycerol 10 %) with protease and phosphatase inhibitors. Yeast whole-cell extracts were prepared by glass bead disruption of cells using a FastPrep system. Protein concentrations were determined and 5 mg of total protein was incubated with 1.5 μl anti-HA-Sepharose beads for 2 hours at 4°C to immunoprecipitate Rpb4/Rpb7-HA. The immunoprecipitates were washed several times with lysis buffer and then incubated with 2 μg of either GST or GST-Ssu72 proteins for 90 min. at 4°C. Reaction mixtures were extensively washed and afterwards electrophoresed on a SDS-polyacrylamide gels, transferred and immunoblotted with the following antibodies: anti-HA, anti-Rpb4 and GST-HRP (A3740, SIGMA).

#### RNA isolation and RT-PCR

Total RNA was extracted with the RNeasy Kit (Qiagen). The cDNA was synthetized with using the iScript™ Advanced cDNA Synthesis Kit (Bio-Rad), following the manufacturer’s instructions. PCR reactions were performed in triplicate with at least three independent cDNA samples.

#### Chromosome Conformation Capture (3C)

3C assay was performed as described in detail in [32].

#### TRO assays

TRO assay was performed essentially as described in [61].

## Acknowledgements

We would like to thank M. Garavís for GST-Ssu72 production, F. Navarro for yeast strains, and M. Hampsey and B. Singh for technical support with 3C assays and providing GST-Ssu72 plasmid. This work was supported by a grant to OC from the Spanish Ministry of Science, Education and Universities (BFU2017-84694-P). OC also thanks to the Program “Escalera de Excelencia” from *Junta de Castilla y León* (CLU-2017-03), co-funded by the P.O. FEDER from *Castilla y León 14-20*. This work was also supported by a grant from National Science Foundation (MCB1020911) and WSU Grant Boost award to AA. ZD was supported by Rumble fellowship from Wayne State University.

## Author contributions

OC conceived the study. PAF, NGP, MJO, BP, ZD, AA, OC performed all experiments. OC and AA designed all experiments, analyzed data, and wrote the manuscript.

## Conflict of interest

The authors declare that they have no conflict of interest

Olga Calvo ORCID # 0000-0002-9786-7916

Athar Ansari ORCID # 0000-0003-2806-421X

## APPENDIX Appendix Supplementary Material

**Table S1.**
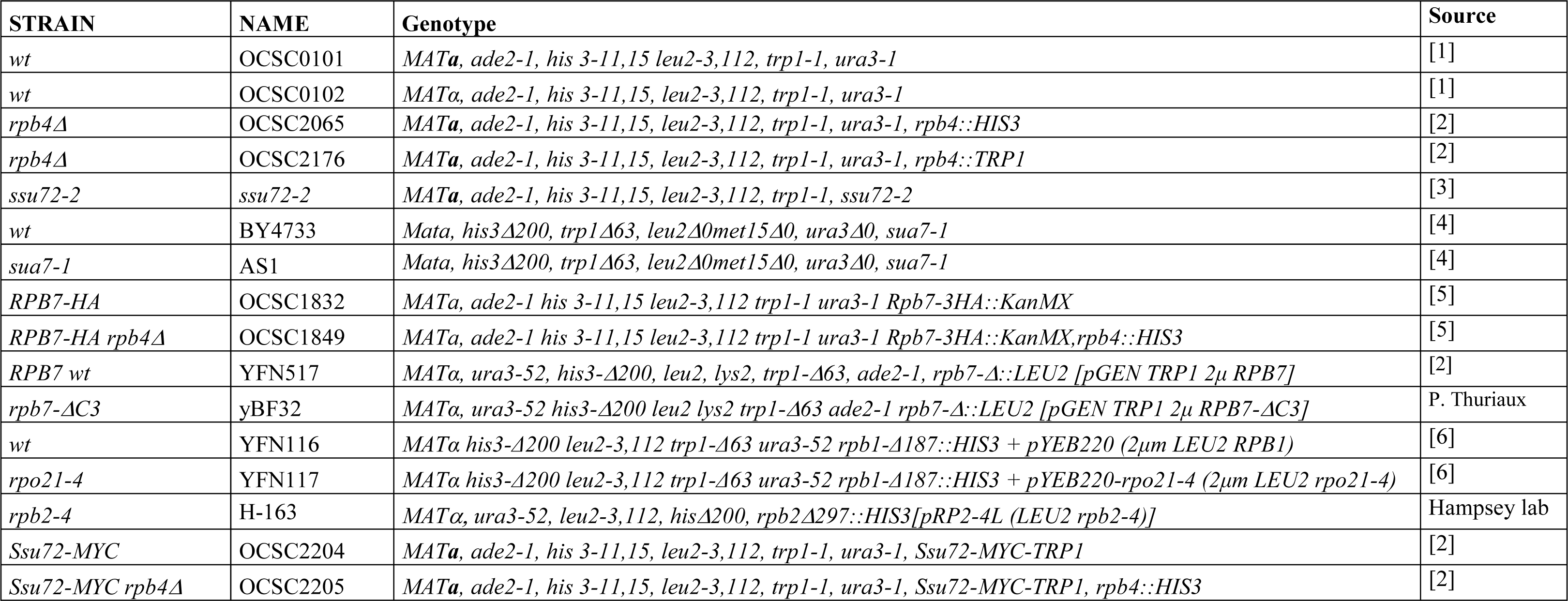
Yeast strains.

**Table S2.** Plasmids.

| Plasmid | Features | Source |
| --- | --- | --- |
| pCM190 | 2μM *URA3* | [7] |
| pCM190-*RPB4* | 2μM *URA3* | [6] |
| pGEN | 2μM *TRP1* | [8] |
| pGEN-*RPB7* | 2μM *TRP1* | [9] |
| pRS423 | *CEN*, *HIS3* | [10] |
| pM647 [pRS423-*SSU72*] | 2μM *HIS3* | [11] |
| pGST-Ssu72 |  | Hampsey Lab |
| pRpb4/Rpb7-6xHis |  | [12] |

**Table S3:**
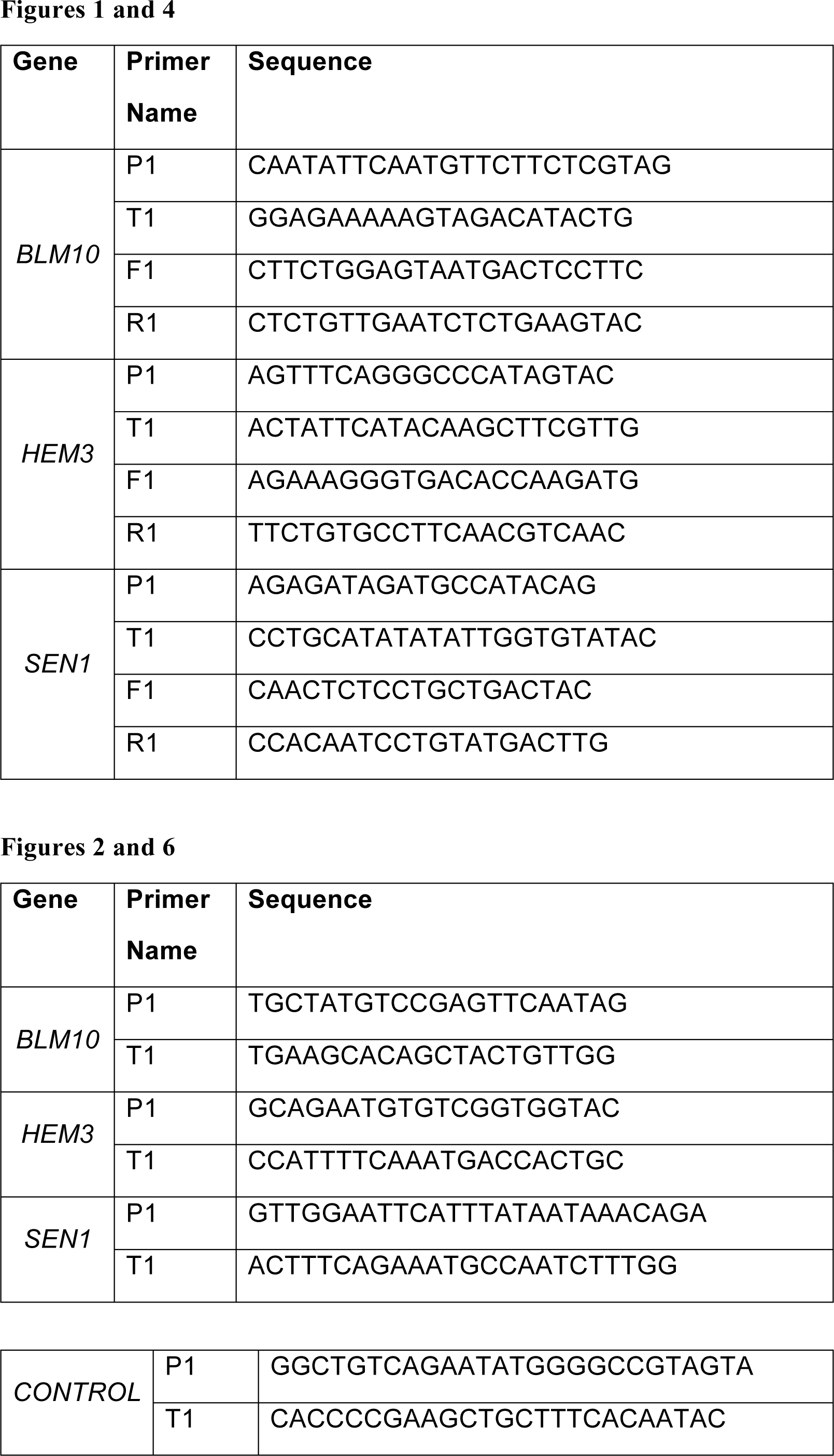
Primers for 3C assay of *BLM10, HEM3* and *SEN1*.

**Table S4:**
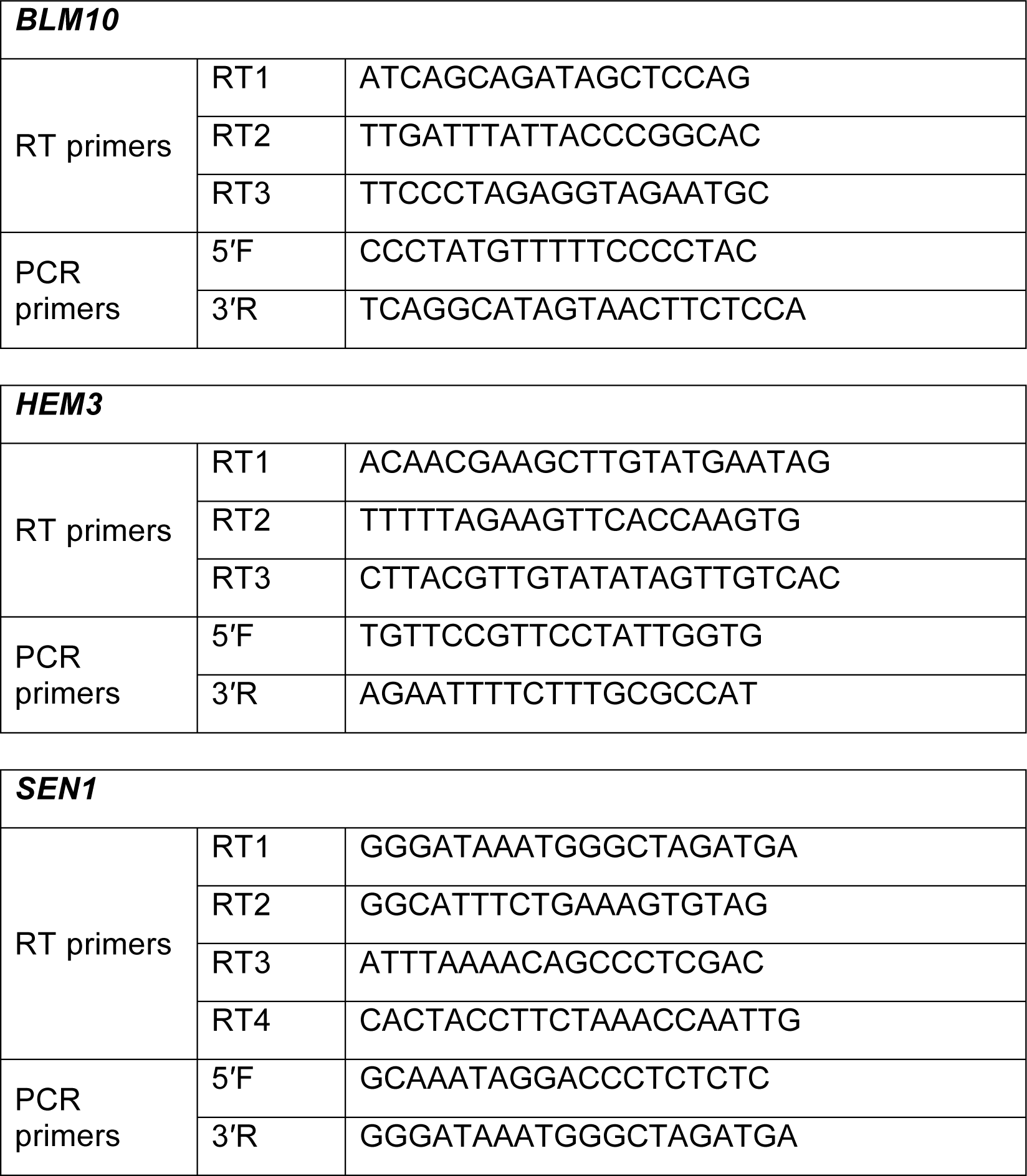
Primers for TRO assay of *BLM10*, *HEM3* and *SEN1*.

